# Direct detection of deformation modes on varying length scales in active biopolymer networks

**DOI:** 10.1101/2023.05.15.540780

**Authors:** Samantha Stam, Margaret L. Gardel, Kimberly L. Weirich

## Abstract

Correlated flows and forces that emerge from active matter orchestrate complex processes such as shape regulation and deformations in biological cells and tissues. The active materials central to cellular mechanics are cytoskeletal networks, where molecular motor activity drives deformations and remodeling. Here, we investigate deformation modes in contractile actin networks driven by the molecular motor myosin II through quantitative fluorescence microscopy. We examine the deformation anisotropy at different length scales in networks of sparsely cross-linked and bundled actin. In sparsely cross-linked networks, we find myosin-dependent biaxial buckling modes across length scales. Interestingly, both long and short-wavelength buckling may contribute to network contractility. In cross-linked bundled networks, uniaxial contraction predominates on long length scales, while the uniaxial or biaxial nature of the deformation depends on bundle microstructure at shorter length scales. The anisotropy of deformations may provide insight to the mechanical origins of contractility in actin networks and regulation of collective behavior in a variety of active materials.

## INTRODUCTION

Active materials are ubiquitous in biology and of increasing interest in engineered materials due to their emergent collective behavior and responsive properties, such as correlated motion, large density fluctuations, and directed force transmission (*1*). These intriguing materials are made of mechanically active building blocks that exert internal forces and produce local motion, resulting in emergent collective phenomena. In biology, active materials emerge from collections of active particles at various length scales, including flocks of birds, suspensions of swarming bacteria, layers of motile cells, and networks of polymers, while engineered active matter includes collections of chemically reactive colloids and mechanically driven particles (*2*). Developing a fundamental understanding of the mechanical properties that emerge from active material microstructure is necessary to characterize functions such as roles within biological tissues and potential for engineered applications such as microrobotics (*3*).

In biological cells, active materials comprise the cytoskeleton, networks and bundles of filament-forming proteins such as actin and microtubules. A primary way that the cytoskeleton is made active is through molecular motors, such as myosin II, that translate the chemical energy of nucleotide hydrolysis into forces that drive filament motion and deformation in actin networks. Myosin II motor heads take force-generating steps, while interacting with multiple actin filaments, resulting in actin filament sliding and buckling that leads to overall network deformation. The activity of motors can also drive relative sliding of filaments and gather filament ends into central locations in a process known as polarity sorting (*4–7*).

These cytoskeletal materials exhibit classic active matter characteristics such as correlated motion (*8, 9*) and large density fluctuations (*8*). However, in contrast to many other active materials, the constituent particles in the cytoskeleton are filaments that are highly deformable and polar, resulting in deformations that depend on orientation and mechanical properties of the filament. For example, actin filaments are semi-flexible polymers, giving rise to an asymmetry in a filament’s response to compression relative to extension. This asymmetric mechanical response leads to actin filaments preferentially buckling under load and is hypothesized to be critical to producing the net contractile behavior characteristic of active cytoskeletal networks (*10–15*). Previous experimental and theoretical studies have suggested that the types of network deformation modes, buckling and sliding, are regulated through the network microstructure and activity such as filament rigidity (*16*), cross-linking between filaments (*13, 16–19*), or motor concentration and properties (*11, 13, 20*). These deformation modes underlie differences in emergent network properties, including whether the material is predominately contractile or extensile. To understand the microscopic origin of the active material behavior, most previous experimental studies have relied on indirect assessment of deformation modes via comparison of contraction rate or other metrics to theoretical models (*17, 18*).

Here, we investigate how active network deformation modes depend on length scale and internal stress through quantitative image analysis of deformation anisotropy (*16*) in cytoskeletal networks with different microstructures. We find that uniaxial deformations, which indicate actin filament sliding (*16*), are the predominate deformation mode in networks of semi-flexible actin at sufficiently low internal stress. At higher myosin densities, the internal stress increases, leading to length-scale dependent biaxial buckling deformations. At intermediate myosin densities, long-wavelength buckling deformations continually rearrange the network persist for tens of minutes without causing network clustering typically observed at higher densities (*12, 21–23*). In networks of cross-linked rigid bundles, by contrast, we observe different patterns of deformation as a function of internal stress and length scale than in networks of filaments. Networks of bundles deform uniaxially on sufficiently large length scales. However, bundles can have different microstructures, where filaments are aligned with the same polarity or with randomly oriented filaments. We find that these different bundle microstructures exhibit either stress-independent (unipolar bundles) or stress and length-scale dependent deformation modes (random polarity bundles). These results reveal how active materials tune deformations at different length scales with microstructure and internal force, emphasizing the importance of considering length scale in building physical models.

## METHODS

### Protein purification, fluorescent labeling, and storage

Actin, skeletal muscle myosin II, filamin, fascin, and fimbrin were purified as described previously and stored as frozen stocks at −80°C (*16, 24, 25*). Actin was purified from rabbit acetone powder (Pel-Freez Biologicals) and stored 2 mM Tris·HCl, pH 8.0, 0.2 mM ATP, 0.2 mM CaCl_2_, 0.2 mM DTT, and 0.005% NaN_3_ (*26*). Myosin II was prepared from fresh chicken skeletal muscle 5 mM PIPES, pH 7.0, 450 mM KCl (*27*). Filamin was purified from chicken gizzard and stored in 10 mM Tris-HCl, pH 7.4, 120 mM NaCl, 0.5 mM EGTA, 1 mM DTT (*28*). Human fascin (GST tag fusion) and pombe fimbrin (HisTag fusion) were purified from E. coli using plasmids from D. Kovar Lab (University of Chicago) (*29, 30*). Actin and myosin II were fluorescently labeled with tetramethylrhodamine-6-maleimide (TMR, Life Technologies) and Alexa-647 maleimide (Life Technologies), respectively (*31*).

### Sample preparation

To prepare microscopy samples, a chamber was constructed from a coverslip passivated with an oil-surfactant layer or lipid bilayer. To prepare an oil-surfacant passivated sample chamber, PFPE-PEG-PFPE surfactant (008; RAN Biotechnologies) was dissolved in a fluorinated oil (Novec-7500 Engineered Fluid; 3M) at a concentration of 2 wt% (*9*). Coverslips (no. 1.5, Fisherbrand) were cleaned by sonication and rinsed in pure water and ethanol. Coverslips were then treated to make hydrophobic by incubating in a 2% vol/vol solution of triethoxy(octyl)silane (440213; Sigma-Aldrich) in isopropanol. The hydrophobic coating was removed from entire coverslip surface except for a 2 mm x 2 mm square region by exposing to UV-ozone for 10 minutes with a square Teflon mask covering the region. This constrained the oil to the square region and prevented flow generated by seeping at the coverslip edges. A glass cylinder was epoxied to the coverslip to produce a microscopy chamber. A minimal volume of oil (~ 3.5 µL) was added to the chamber to coat the surface, and excess solution removed immediately prior to adding the actin polymerization mixture.

To prepare a lipid bilayer passivated sample chamber (*32*), coverslips (no. 1.5; Fisherbrand) were first rinsed with water and ethanol then cleaned through exposure to UV-ozone for 20 min. A glass cylinder was epoxied to the clean coverslip, and vesicle buffer (10 mM sodium phosphate, pH 7.5, and 140 mM NaCl) added as soon as possible. DOPC vesicles (1,2-dioleoyl-sn-glycero-3-phosphocholine; Avanti Polar Lipids) were then added to a concentration of 100 μM. The coverslip was incubated with the vesicles for 15 min to allow bilayer formation, before the bilayer was rinsed with a buffer comprising of 1x F-buffer (below, in actin polymerization) prior to adding the final sample.

#### Sparsely cross-linked actin networks

To prepare cross-linked actin networks crowded to a coverslip surface (*12*), a mixture of 0.3 wt% 15 cp methylcellulose, 1 µM actin (1:10 TMR-maleimide labeled:unlabeled), 0.002 µM filamin crosslinker, and an oxygen scavenging system to prevent photobleaching [4.5 mg/mL glucose, 2.7 mg/mL glucose oxidase (345486; Calbiochem), 1,700 units/mL catalase (02071; Sigma), and 0.5 vol/vol % β-mercaptoethanol], was made in 1x F-buffer (10 mM imidazole, pH 7.5, 1 mM MgCl2, 50 mM KCl, 0.2 mM EGTA, and 4 mM ATP). Actin was allowed to polymerize and crowd to the coverslip surface for 30 minutes before addition of dimeric Alexa-647 labeled myosin.

#### Bundled actin networks

Bundled actin networks were constructed similarly to sparsely cross-linked networks, but have no cross-linker (filamin) during the initial 30 min polymerization period. To prepare fascin cross-linked bundles, fascin was added at a ratio of 1:10 fascin:actin to the polymerized actin sample and incubated for 20 minutes to allow bundle formation. For additional filamin cross-linking, filamin was subsequently added at a ratio of 1:500 filamin:actin and incubated for an additional 20 minutes. For fimbrin bundles, the fimbrin was added 20 minutes after actin polymerization at a ratio of 1:10 and incubated for 20 minutes. Myosin filaments were polymerized separately for 10 minutes in the same buffer conditions described above for actin polymerization before addition to the bundled actin networks.

### Image acquisition and analysis

An inverted microscope (Eclipse Ti-E; Nikon) with a spinning-disk confocal head (CSU-X; Yokagawa Electric) was used for all imaging. Imaging was done under magnification of a 40× 1.15 N.A. water-immersion objective (Apo LWD; Nikon) or a 60×/1.49 NA oil immersion objective (Zeiss). Images were acquired with a CMOS camera (Zyla-4.2-USB3; Andor) or a CCD camera (Coolsnap HQ2, Photometrics). Images were acquired every 1 s for Figs. 1–5 and every 5 s for Fig. 6. Calculation of flow vectors and deformation anisotropy was conducted as we described previously (*16*) using PIV software (*33*) and custom Matlab scripts. Values of *ρ*_myo_ and *P*_biaxial_ were smoothed such that data points report the average value over overlapping windows of 50 s in fimbrin-bundled sample and 20 s in all other samples.

**Figure 1:**
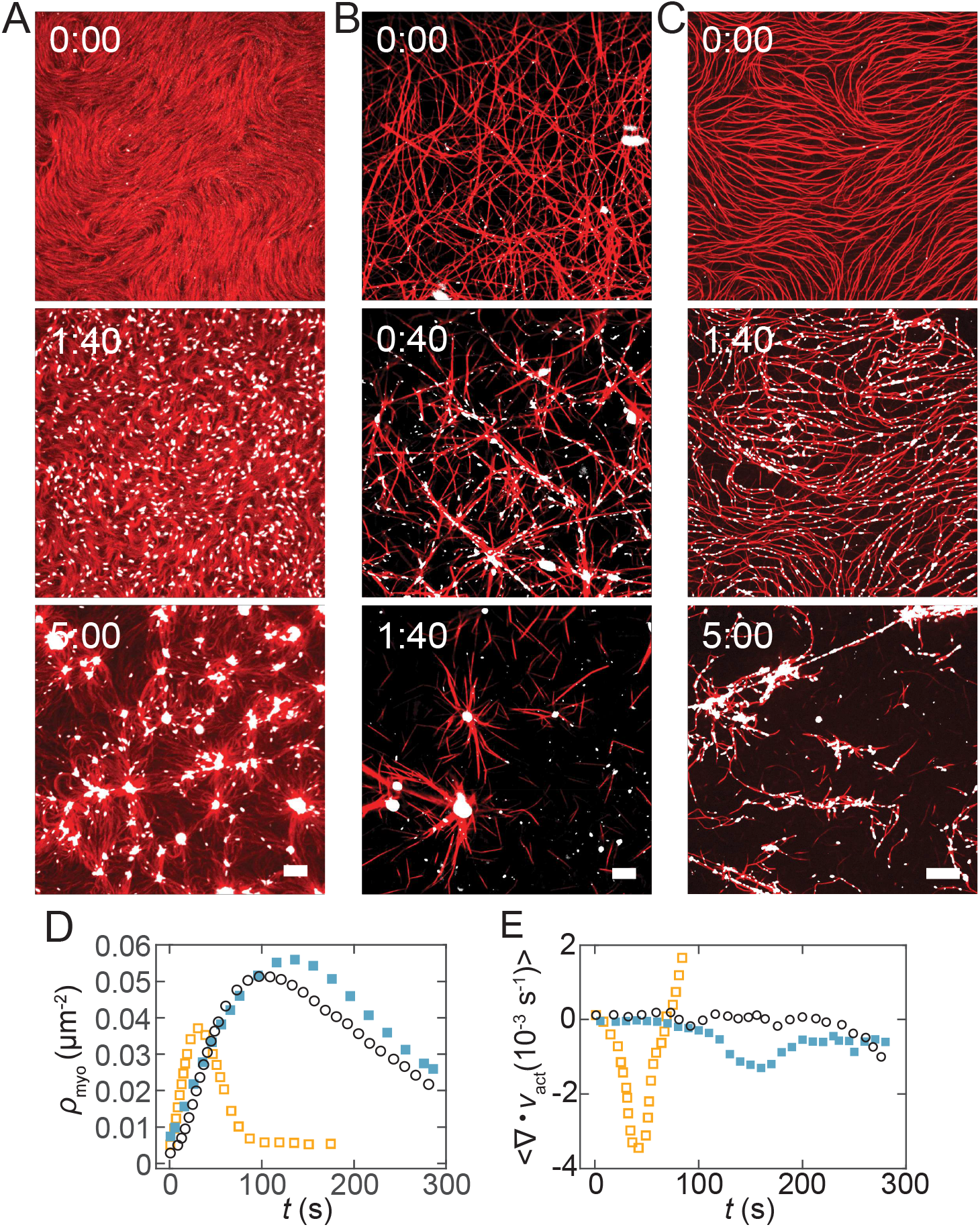
Actomyosin network contractility is dependent on filament cross-linking and bundling. (A) Fluorescence microscopy images of a network of unbundled actin filaments (red) sparsely cross-linked with the flexible cross-linker filamin as myosin (white) accumulates over time. (B) Images of unipolar actin bundles (red) formed with a rigid actin cross-linker (fascin) and then linked together with the flexible cross-linker filamin as myosin (white) accumulates. (C) Images of actin bundles (red) with random actin polarity formed by the cross-linker fimbrin as myosin (white) accumulates. In each image sequence, the myosin is added to the sample at time 0:00 and subsequently polymerizes into punctate filaments. Scale bars are 10 µm. (D) Myosin filament puncta density, *ρ*_myo_, plotted against time after myosin addition, for the sparsely cross-linked (black open circles), unipolar bundles (orange open squares) and random polarity bundle (filled blue squares) samples. (E) Divergence as a function of time for representative samples of sparsely cross-linked (open black circles), unipolar bundle (open orange squares), and random polarity bundle (filled blue squares) actin networks.

### RESULTS

To create a cytoskeletal material inspired by the cellular cortex, we crowd actin filaments to a thin, dense layer with a depletion agent (see Methods) (*12*). Filamin, an actin cross-linking protein, connects entangled filaments into networks without causing bundling when incorporated at a low concentration (0.01 µM; 1:100 molar ratio with respect to actin monomer). A thin oil-surfactant layer passivates the coverslip surface against protein adhesion (*9*). To introduce activity, we add myosin II dimers. We define the time the motors are first added to the sample as time *t* = 0 s. Upon dilution in the sample buffer, which contains lower salt concentration than the storage buffer, the myosin II dimers polymerize into puncta (referred to as myosin, Fig. 1, white puncta), which are filaments that each contain several hundred myosin II motors (*12*).

Myosin incorporates into the sparsely cross-linked network, initially increasing from 0 to 0.05 puncta/µm^2^ over the first ~100 s, as myosin polymerizes into elongated puncta that bind to actin filaments (*ρ*_myo_, Fig. 1D, open circles). During this initial period of myosin accumulation, there is an increase in the fluctuations along the contour of actin filaments, indicating myosin-induced stresses bending actin filaments. This filament bending induces local filament rearrangement without substantially changing network density (Fig. 1A, 0:00-1:40). As myosin continues to accumulate, the actin and myosin filaments rearrange into clusters with a central myosin core and intervening voids (Fig. 1A, 5:00). This is consistent with previous reports of actomyosin network coarsening *in vitro*, where dense actomyosin clusters form given a sufficiently high myosin concentration (*12, 21–23*). As a result of clustering, the number density of myosin puncta begins to decrease near *t* = 100 s (Fig. 1D).

We previously found that cross-linked networks of unipolar rigid bundles contract via uniaxial deformations arising from relative sliding of the bundles (*16*). These networks are constructed with two different actin cross-linkers. One type of cross-linker, fascin, rigidly links actin filaments together into bundles with a uniform polarity (*34*). We observe that these bundles appear as straight rods with diffraction limited width that may overlap with other bundles (Fig. 1B, (*16, 24*)). We then add a flexible cross-linker, filamin, after the fascin-mediated bundles form to ensure that the bundles form an inter-connected network. Effectively, the unipolar bundles act as polar filaments with a higher stiffness than the unbundled, semi-flexible actin (*24*). The unipolar bundle networks form asters upon addition of myosin (Fig. 1B, 1:40, (*16*)). As with the sparsely cross-linked network in Fig. 1A, the density of myosin puncta initially increases as myosin accumulates in the network, and then decreases as clusters form (Fig 1D, open squares).

To more broadly understand the influence of actin bundle structure on network deformation modes, we use a different rigid actin cross-linker, fimbrin, to make actin bundles. These bundles have similar packing of the actin filaments as fascin bundles (*35*), and therefore might also contract uniaxially. In contrast to fascin bundles, within fimbrin bundles are aligned with random polarity (*36*). Myosin filaments can simultaneously bind to actin filaments with opposite polarity and build internal stress within a bundle (*24*).

Myosin activity induces the random polarity bundle network to coarsen into dense actomyosin clusters (Fig. 1C), similar to the unipolar bundle network. However, the intermediate stages of network reorganization are distinct from the unipolar bundle network. While the unipolar bundles appear rigid, there is visual evidence of buckling or bending of the random polarity bundles (Fig. 1C, 1:40). Similar to the previous networks, the myosin puncta density initially rises and then falls during network clustering (Fig 1D, filled squares (*16*)).

The average divergence of actin displacement vectors calculated by particle imaging velocimetry (PIV, *Materials and Methods*) has been used previously to quantify contractility in actomyosin networks (*20*). The typical behavior of the average divergence over time is a decrease to negative (contractile) values as myosin accumulates and contracts the network followed by a return to near zero values as the network forms clusters (*16, 20*). The bundled networks from Fig. 1B and 1C both display this expected behavior with minimum values of the divergence (Fig. 1E, orange open squares and blue closed squares) reached near the respective myosin puncta density peaks in Fig. 1D. However, the average divergence of the sparsely cross-linked sample oscillates near zero throughout most of the network rearrangement and only decreases to consistently negative values long after network clustering begins (Fig. 1E, black open circles). This is surprising given that the buckling/compression of filaments observed in Fig. 1A is expected to produce a negative divergence (*16, 20*).

To obtain a better understanding of differences in the contraction of these networks, we develop a metric for contraction on different length scales. We split the actin images into boxes of size *s* which contain local PIV vectors, 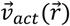 (Fig. 2A). The contraction of each box may be characterized by the contractile moment:

**Figure 2:**
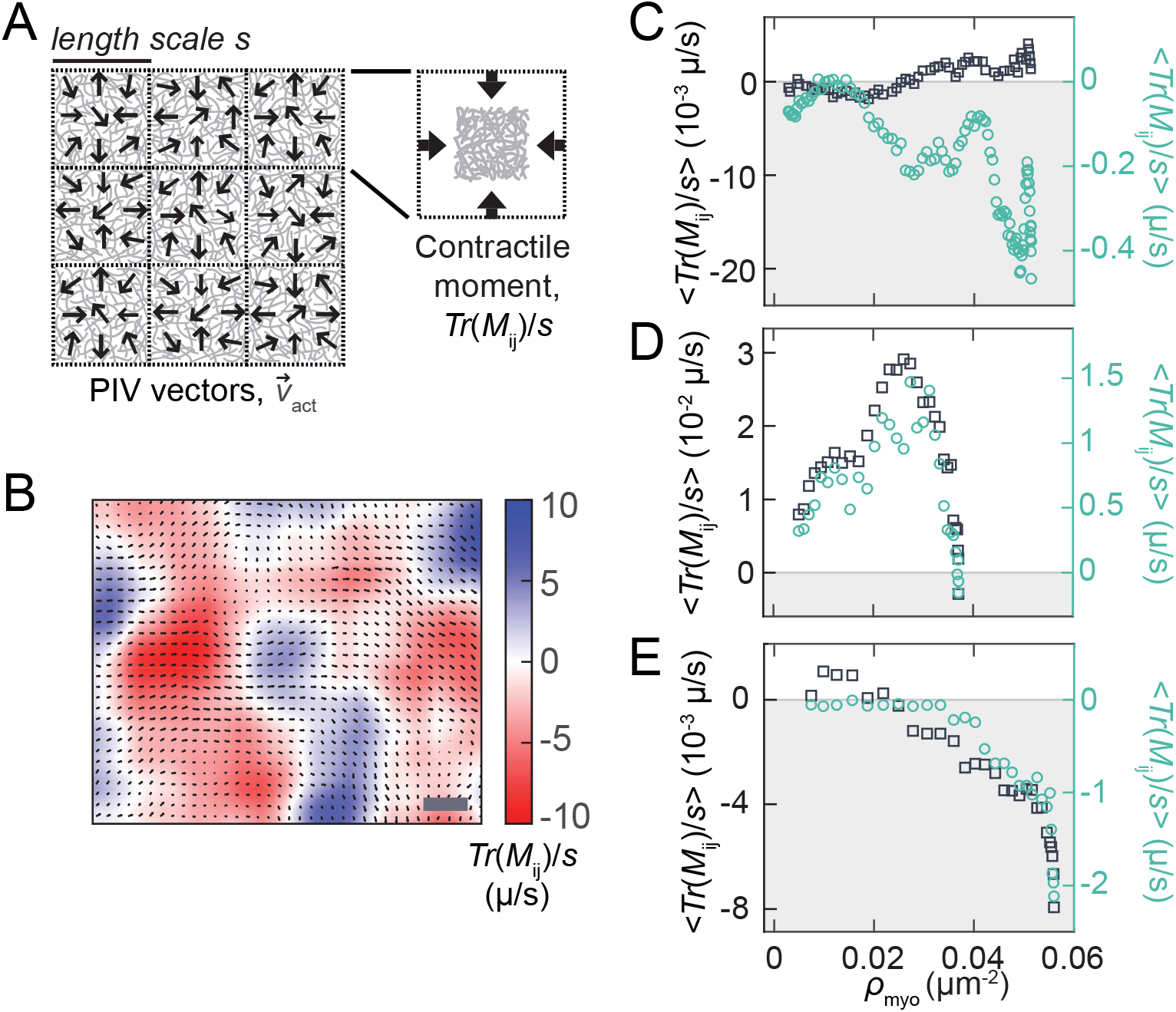
Contractility is dependent on actin network architecture and length scale. (A): Particle imaging velocimetry generates vectors describing the local material flow in an actin network. The network images are separated into boxes of side length *s*, and the contractility of each box is quantified using the contractile moment. (B): Contractile moment values of overlapping boxes in the sparsely cross-linked network during contraction. The color represents Tr(*M*_ij_)/*s* of a box with *s* = 50 µm centered at each position. (C): Spatial average of Tr(*M*_ij_)/*s* in non-overlapping boxes with *s* = 50 µm (open green circles) and *s* = 10 µm (open black squares) in sparsely cross-linked network. (D): Spatial average of Tr(*M*_ij_)/*s* in non-overlapping boxes with *s* = 50 µm (open green circles) and *s* = 10 µm (open black squares) in unipolar bundle network. (E): Spatial average of Tr(*M*_ij_)/*s* in non-overlapping boxes with *s* = 45 µm (open green circles) and *s* = 6.5 µm (open black squares) in random polarity bundle network.

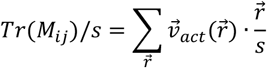

where 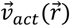 is the PIV vector located at position 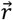 from the center of the box and *M*_ij_ is the flow dipole moment tensor (*16*). This has been rescaled from our previous quantification of contraction (*16*) to facilitate comparison of contractile moments with different *s* (length scale) values.

At a sufficiently large length scale, a decrease of <Tr(*M*_ij_)/*s*> to negative (contractile) values during the initial stages of contraction occurs in the sparsely cross-linked network (Fig. S1A). To explain the trends revealed by the average contractile moment, we visualize contraction on mesoscopic length scales by calculating Tr(*M*_ij_)/*s* with *s* = 50 µm for overlapping boxes and plotting the value using the color scales in Fig. 2B and Fig. S2. In all networks, large areas with contractile (red) Tr(*M*_ij_)/*s* values are separated by regions of extensile (blue) Tr(*M*_ij_)/*s* values. Interestingly, these contractile regions begin developing at early time points and low myosin densities and increase in intensity as the contraction proceeds (Fig. S2).

Given that the deformation modes of these networks are expected to vary between filament buckling and sliding (*16*), we anticipate that the contractility on different length scales will depend on increasing internal stress in divergent ways. To examine this possibility, we plot the average contractile moment in non-overlapping boxes against *ρ*_myo_ during the myosin accumulation stage (as defined Fig. 1D). In the sparsely cross-linked network, <Tr(*M*_ij_)/*s*> at *s*=50 µm begins decreasing to negative values near *ρ*_myo_ =0.01 µm^−2^ before oscillating slightly between *ρ*_myo_=0.02 µm^−2^ and 0.04 µm^−2^ (Fig. 2C, green open circles). A steeper decrease then occurs near *ρ*_myo_=0.04 µm^−2^ followed by a return to somewhat less negative values at *ρ*_myo_=0.05 µm^−2^. The magnitude of <Tr(*M*_ij_)/*s*> at *s*=10 µm is smaller and increases to slightly positive values at a similar *ρ*_myo_ as the decrease in <Tr(*M*_ij_)/*s*> at *s*=50 µm (Fig. 2C, open black squares).

The changes of <Tr(*M*_ij_)/*s*> with increasing *ρ*_myo_ differ in the bundled networks. In the unipolar bundle network, the magnitude of <Tr(*M*_ij_)/*s*> is also smaller for the shorter length scale, *s*=10 µm, than *s*=50 µm (Fig. 2D). However, the trends of the curves at both length scales are similar, with both starting at initially positive values before decreasing to negative ones. This decrease continues after the peak in myosin density (Fig. S1B).

Similarly, the curves for <Tr(*M*_ij_)/*s*> versus *ρ*_myo_ at longer and shorter length scales in the random polarity bundled network have different magnitudes but otherwise resemble one another closely (Fig. 2E, *s*=6.5 µm, open black squares and *s*=45 µm, open green circles). In this network, both remain near zero until they decrease to negative values when *ρ*_myo_ exceeds approximately 0.03 µm_-2_. Overall, the trends of contractility with internal stress and length scale are very different between these networks.

To characterize the origin of contractility in terms of filament sliding versus buckling deformation modes, we diagonalize the flow dipole moment tensor, *M*_ij_, of boxes with varying *s* (*16*). The resulting eigenvalues, *M*_min_ and *M*_max_, characterize the local deformation anisotropy (Fig. 3A). A value of *M*_min_/*M*_max_ of 0 indicates that the deformation of actin network within the defined box is along a single axis. At the other extreme, a value of *M*_min_/*M*_max_ of 1 indicates that the deformation is completely biaxial. We superimpose ellipses on the actin images with minor and major axes *M*_min_ and *M*_max_ respectively to visualize the anisotropy of deformations. For *s* = 20 µm in the sparsely cross-linked network, the size of the ellipses (magnitudes of *M*_min_ and *M*_max_) increases between time 0:00 to 1:40 (Fig. 3B), indicating an increase in extent of deformation as myosin accumulates in the network. At a smaller length scale, *s* = 5 µm, we also find that *M*_min_ and *M*_max_ increase as myosin accumulates (Fig 3C).

**Figure 3:**
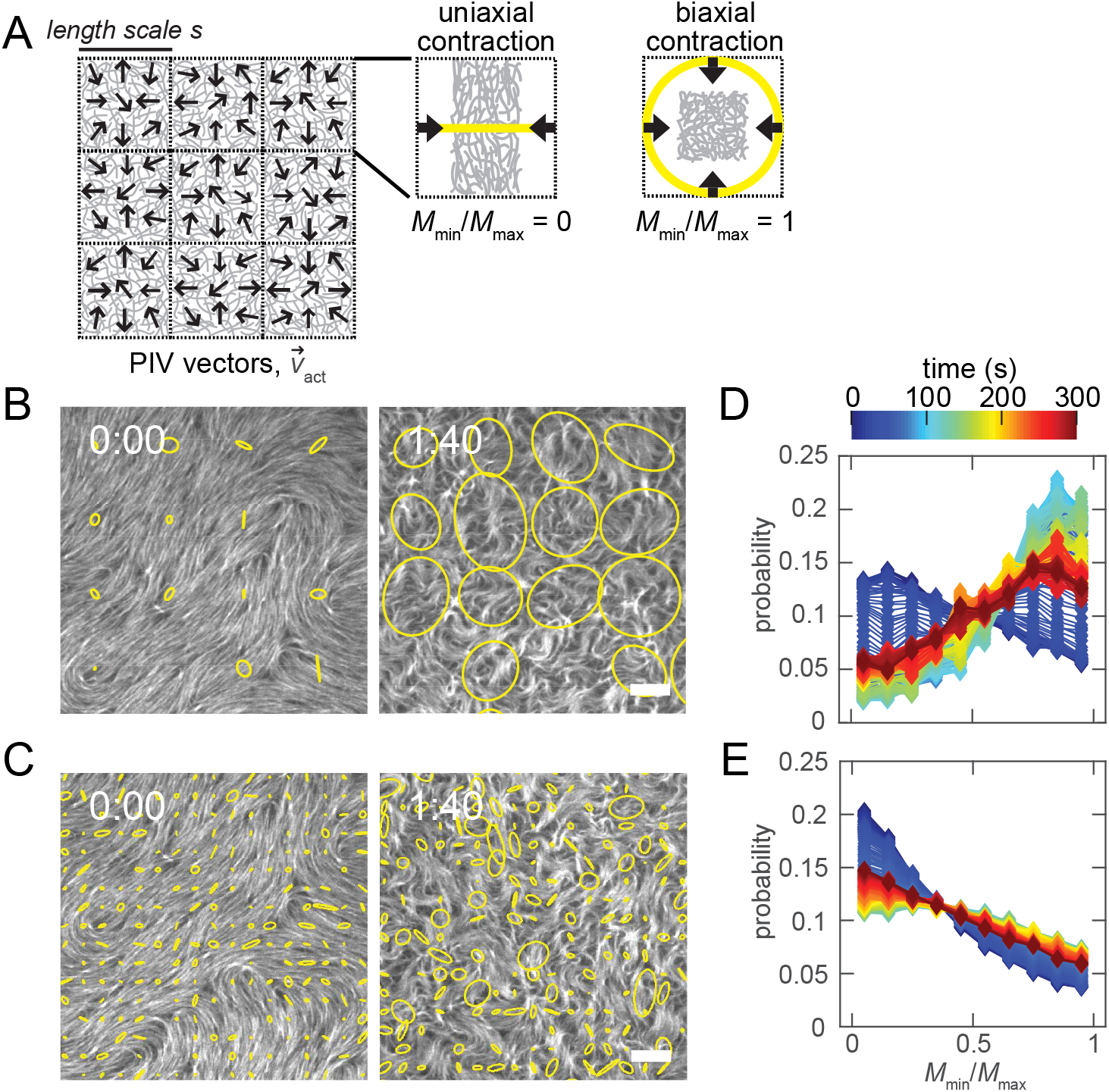
Anisotropy analysis reveals varying deformation mechanisms in contractile networks. (A): The deformation anisotropy of each box of PIV vectors is quantified using the flow dipole tensor and represented by an ellipse. (B): Deformation anisotropy ellipse images in sparsely cross-linked network at length scale *s* = 20 µm. (C): Deformation anisotropy ellipse images in same network at *s* = 5 µm. (D): Distributions of the deformation anisotropy, *M*_min_/*M*_max_, at varying time points with *s* = 20 µm. (E): Distributions of *M*_min_/*M*_max_ with *s* = 5 µm. Scale bars in (B) and (D) are 10 µm.

The deformation anisotropy also changes as the sparsely cross-linked network reorganizes. Deformations are predominantly uniaxial when myosin density in the network is initially low (left panels in Fig. 3B and 3C, quantified in Fig. 3D and 3E, blue curves). This differs from our previously reported data of networks with greater initial myosin density, which exhibited biaxial deformations arising from filament buckling (*16*). As network deformation proceeds, corresponding to increased myosin accumulation, we observe a large shift in the probability distribution of *M*_min_/*M*_max_ at *s* = 20 µm. The probability of biaxial deformations (*M*_min_/*M*_max_>0.5) increases such that the distribution becomes peaked at *M*_min_/*M*_max_ ~ 0.8 (Fig. 3D). At a smaller length scale (*s* = 5 µm, Fig. 3E), the distribution also shifts to more biaxial deformations after the addition of myosin, but distribution of *M*_min_/*M*_max_ always remains more uniaxial and the changes over time are smaller. At long times on both length scales, the distribution eventually shifts back to a distribution with fewer biaxial deformations.

To understand how these patterns of deformation give rise to the differences in contractility seen in Fig. 2, we define *P*_biaxial_ as the proportion of deformations in the network at a given time and *s* value with *M*_min_/*M*_max_ values greater than 0.5. We then plot *P*_biaxial_ as a function of *ρ*_myo_ during the myosin accumulation stage similarly to the plots of <Tr(*M*_ij_)/*s*> in Figs. 2C-2E. We find that *P*_biaxial_ with *s* = 5-10 µm (Fig. 4A, open black squares) is consistently lower than *P*_biaxial_ with *s*=45-50 µm (Fig. 4A, open green circles) in the sparsely cross-linked network at all values of *ρ*_myo_. At both length scales, *P*_biaxial_ is initially constant over a range of low *ρ*_myo_, but sharply increases above a threshold *ρ*_myo_. The increase in *P*_biaxial_(*s*) at s = 5-10 µm occurs at a higher value of *ρ*_myo_, which is consistent with a higher internal stress requirement for small wavelength buckling modes. Also consistent with this stress requirement, the value of *ρ*_myo_ required to raise *P*_biaxial_ by 20% of its baseline value (*ρ*_activate_) decreases with *s* in an average of four samples (Fig. S3).

**Figure 4:**
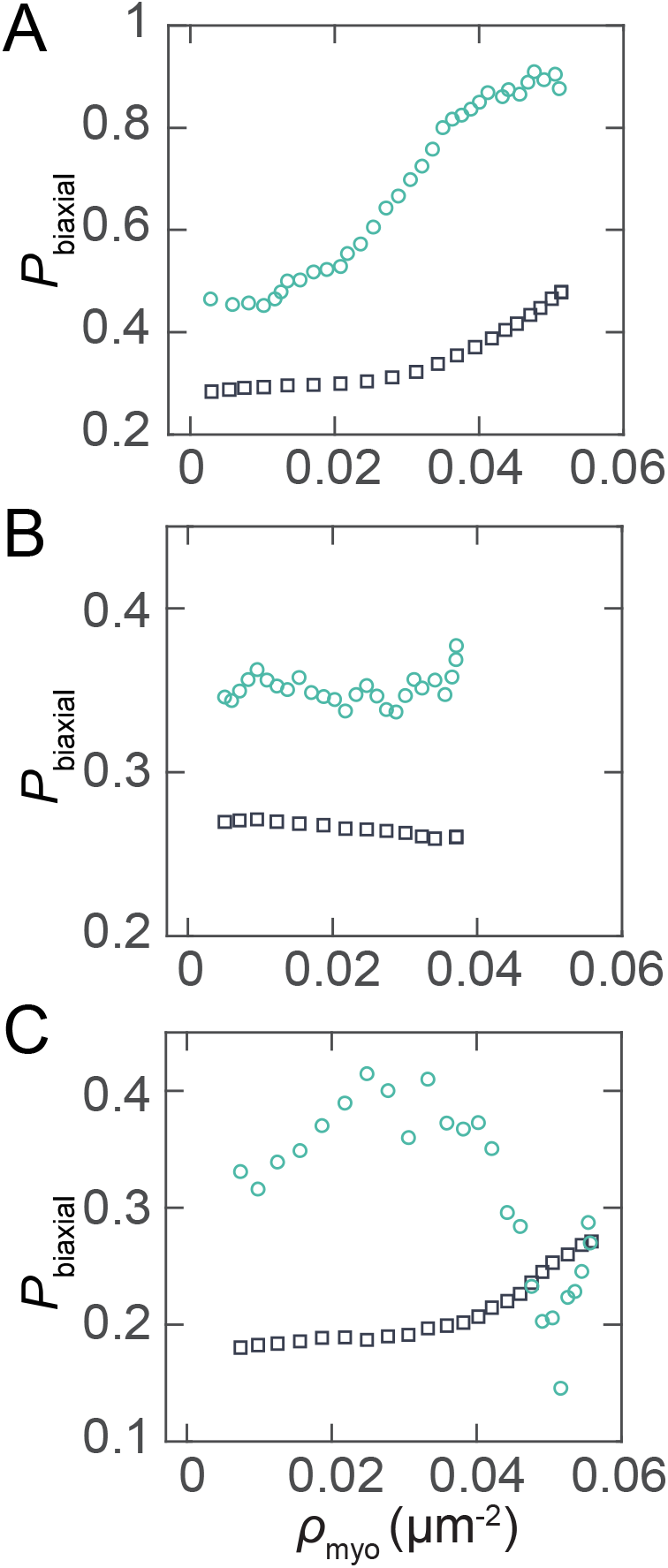
Length-scale dependent activation of deformation modes varies for different actin networks. *P*_biaxial_(*s*), defined as the fraction of *M*_min_/*M*_max_ values greater than 0.5 as a function of myosin puncta density during the myosin accumulation stage for (A) sparsely cross-linked networks, with *s* = 5-10 µm (open black squares) and *s* = 45-50 µm (open green circles); (B) unipolar bundle networks, with *s* = 5-10 µm (open black squares) and *s* = 45-50 µm (open green circles); and (C) random polarity bundle networks, with *s* = 3-7 µm (open black squares) and *s* = 42-46 µm (open green circles).

Intriguingly, the sharp increase of the long-wavelength biaxial buckling modes in Fig. 4A occurs at approximately the same *ρ*_myo_ value as the initial decrease of <Tr(*M*_ij_)/*s*> with *s*=50 µm in Fig. 2C (*ρ*_myo_ ≈ 0.02 µm^−2^). This suggests that long-wavelength buckling starts contributing to contractility before the short-wavelength buckling begins. The additional decrease of <Tr(*M*_ij_)/*s*> with *s*=50 µm at *ρ*_myo_ ≈ 0.04 µm^−2^ occurs while *P*_biaxial_ (*s*=5-10 µm) is increasing and *P*_biaxial_ (*s*=45-50 µm) is reaching a plateau. However, the value of <Tr(*M*_ij_)/*s*> at *s*=10 µm in Fig. 2C remains slightly extensile even during the increase of *P*_biaxial_ at the shorter length scale.

The unipolar bundle network (Fig. 4B) similarly has a baseline value of *P*_biaxial_ that is lower for *s* = 5-10 µm (open black squares) than *s* = 45-50 µm (open green circles). Unlike the sparsely cross-linked network in Fig. 4A however, there is no sharp increase in *P*_biaxial_ with increasing *ρ*_myo_. The difference of *P*_biaxial_ from its initial value to its extremum at or before the peak in *ρ*_myo_, Δ*P*_biaxial,_ remains similarly near zero over a range of *s* (Fig. S4). Therefore, the uniaxial sliding deformations underlying contractility in this network (*16*) dominate the deformation anisotropy at both length scales.

For the random polarity bundle network, a different pattern of length-scale dependent changes in *P*_biaxial_ with increasing myosin density is observed (Fig. 4C). At a smaller value of *s*=3-7 µm (open black squares), *P*_biaxial_ begins increasing at *ρ*_myo_ ≈ 0.04 µm^−2^ while at *s*=42-46 (open green circles), *P*_biaxial_ decreases. This suggests that buckling of bundles occurs locally, but the bundles remain sufficiently intact to transmit anisotropic deformations on longer length scales, which is consistent with the images in Fig. 1C. The value of Δ*P*_biaxial_ crosses over from positive to negative values at s > 25 µm (Fig. S4).

Previous studies have suggested that contraction may originate at the microscale in proximity to myosin filaments and propagate outward into the network (*15, 19*). However, the sparsely cross-linked network appears to contrast with this expectation given that the onset of buckling with increasing *ρ*_myo_ first occurs at higher *s*. To specifically inhibit the buckling modes at small *s* and examine the influence on contractility, we investigate networks with lower myosin densities. In a sparsely cross-linked networks with ~ 5 times lower *ρ*_myo_ than the network in Fig. 1A (Fig. 5A, filled squares), we observe no coarsening of the actin network even after 15 minutes of myosin activity (Fig. 5B). When we investigate network with a slightly increased *ρ*_myo_, (Fig. 5A, open circles), we observe a restoration of some actin coarsening at *t*=15:00. (Fig. 5C). Clustering of the myosin occurs in both networks as indicated by the decreases in *ρ*_myo_ following their peaks in Fig. 5A. Large regions with contractile Tr(*M*_ij_)/*s* values are present in both samples (Fig 5D, red areas) but are lower magnitude than the data in Fig. 2B.

**Figure 5:**
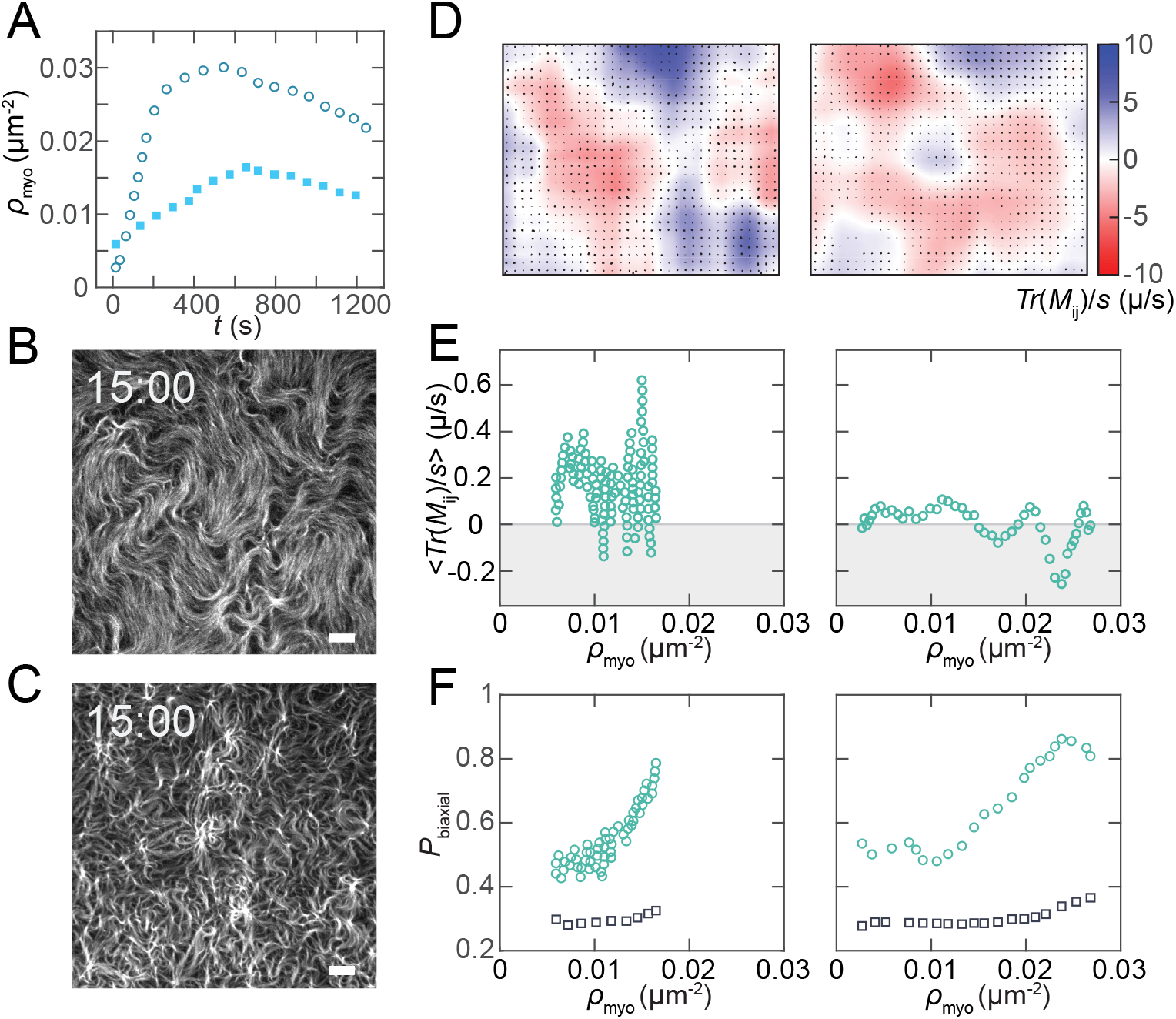
Reducing the myosin density in sparsely cross-linked actin networks lowers the frequency and net contractility of buckling. (A) Myosin filament puncta density, *ρ*_myo_, plotted against time after myosin addition in two sparsely cross-linked networks with low *ρ*_myo_ (open circles) and intermediate *ρ*_myo_ (filled squares). (B) Image of actin in low *ρ*_myo_ network after 15:00 of myosin activity. (C) Image of actin in intermediate *ρ*_myo_ network after 15:00 of myosin activity. (D) Heat maps of Tr(*M*_ij_)/*s* in overlapping, 50 µm boxes in low *ρ*_myo_ (left) and intermediate *ρ*_myo_ (right) networks. (E) Spatial average of Tr(*M*_ij_)/*s* with non-overlapping boxes, s = 50 µm, in low *ρ*_myo_ (left) and intermediate *ρ*_myo_ (right) networks. (F) *P*_biaxial_ with *s* = 45-50 µm (green circles) and *s* = 5-10 µm (black squares) in low *ρ*_myo_ (left) and intermediate *ρ*_myo_ (right) networks.

We repeat our analysis of <Tr(*M*_ij_)/*s*> with s=50 µm in these samples to assess how contractility varies with deformation mode differences. In the sample with the lowest *ρ*_myo_, <Tr(*M*_ij_)/*s*> (*s*=50 µm) remains mostly extensile and oscillates with increasing *ρ*_myo_ (Fig. 5E, left panel). This contrasts with the decrease to negative values seen in Fig. 2C. In the intermediate *ρ*_myo_ sample, the decrease to negative <Tr(*M*_ij_)/*s*> with increasing *ρ*_myo_ is qualitatively restored (Fig. 5E, right panel), though the minimum value is somewhat less negative than Fig. 2C. Interestingly, the small decrease between *ρ*_myo_=0.01 µm^−2^ and *ρ*_myo_=0.02 µm^−2^ and the subsequent steeper decrease at *ρ*_myo_>0.02 µm^−2^ resembles the trend in Fig. 2C.

Examining *P*_biaxial_ during these changes in <Tr(*M*_ij_)/*s*> reveals potential contributions of individual deformation modes to contractility. In the lowest *ρ*_myo_ network, *P*_biaxial_ (*s*=45-50 µm) sharply increases near *ρ*_myo_ = 0.01 µm^−2^ while only a small increase in *P*_biaxial_ (*s*=5-10 µm) occurs (Fig. 5F, left panel). In the intermediate *ρ*_myo_ network, the trends of both *P*_biaxial_ curves are initially similar, but both reach slightly higher peak values (Fig. 5F, right panel). Similar to the high *ρ*_myo_ network in Fig. 2C and 4A, the initial decrease of <Tr(*M*_ij_)/*s*> in the intermediate *ρ*_myo_ network occurs during the increase in *P*_biaxial_ (*s*=45-50 µm) between *ρ*_myo_ = 0.01 µm^−2^ and *ρ*_myo_ =0.02 µm^−2^ (Fig. 5E and Fig. 5F, right panels). This suggests that long wavelength buckles are again contributing to contractility. However, <Tr(*M*_ij_)/*s*> does not decrease during the rise of *P*_biaxial_ (*s*=45-50 µm) in the low *ρ*_myo_ network (Fig. 5E and Fig. 5F, left panels). Similar maximum values of *P*_biaxial_ (*s*=45-50 µm) happen in the low *ρ*_myo_ network compared to the more contractile networks, but they require less myosin and are reached at ~700 s versus ~100 s after myosin addition (Fig. S5). Thus, buckling at low *ρ*_myo_ might be mechanically different and less contractile and/or contractile buckling is counterbalanced by extensile deformations.

## DISCUSSION

Deformations in active, soft materials are critical during processes requiring changes of material shape, size, or transport of components. The mode of deformation at the microscale can include both translocation of components with respect to one another and deformation of components, regulating material scale mechanical responses such as contraction or expansion (*13*), flow-induced transport (*9, 37*), or force transmission to external physical attachments such as cellular adhesions. To understand how microscopic material structure influences deformations and mechanical response, substantial efforts have been made connecting theoretical and computational models of material deformation to available experimentally observable quantities (*38*). Here, we build on our previous work in which deformation anisotropy was analyzed at a single length scale to identify actin buckling and sliding (*16*) to bridge the gap between experiments in which deformation modes typically must be inferred from indirect measurements and modeling in which the deformations on all length scales are directly accessible. For example, our analysis may be useful for testing models for how energy is injected and dissipated on varying length scales in active matter systems (*39*). While models describing contraction as propagating out from a local buckling deformation and being amplified by the network (*14, 40*) may be appropriate for some contracting materials, to our knowledge there is currently no model predicting whether buckling can cascade to smaller length scales. Such a possibility may be consistent with the sparsely cross-linked network deformations we observe here and is an interesting topic for future studies.

Analysis of deformation modes may also clarify contraction mechanisms in the more complex, composite cytoskeletal materials inside of living cells. Intracellular actin networks include disordered meshworks, branched networks that may inhibit contractility (*41, 42*), and bundles that may regulate myosin-II force magnitudes (*24*) and transmit contractile forces to other actin structures (*43*). As a result of these actin architectures, multiple deformations modes could exist within the same cell. Strategies to label actin and detect flow in cells (*44*) are likely adaptable to linking specific modes with contractility similarly to the analysis performed here. Downstream analysis could examine co-regulatory roles of actin binding proteins and specific contractile deformations. For example, small versus large-wavelength buckling could lead to filament severing (*12, 15*) and affect the activity of actin polymerizing proteins that act on the generated filament barbed ends. Force transmission also occurs between different filaments of the cytoskeleton, with microtubules potentially becoming bent or broken in response to contractile forces generated by actin networks (*45, 46*). Additionally, given the presence of feedback between localized signaling and contractility (*47–49*), future studies could also establish actin deformations as indicators of proximal signaling dynamics.

Force transmission may also occur in organelles outside of the cytoskeleton, and analysis of the resulting deformations could be used as a step toward understanding when, how, and why forces are transmitted in these active materials. For example, recent research has indicated that organelle-organelle contacts are tethers and potentially capable of exerting forces to divide mitochondria (*50, 51*). This suggests that force transmission occurring at these sites may be important for maintaining normal organelle structure and distribution. Altering the physical structure of mitochondria via deformation and translocation may influence their metabolism (*52*) and, consequently, their ability to produce energy or induce detrimental effects that lead to cell aging or death. As another example, mechanosensitive processes such as the opening of ion channels may be related to deformations in both cells and the fibrous networks of the extracellular matrix (*53*). More generally, analysis of deformation modes in cells could function as an accessory to omics-type approaches that help link individual or sets of molecules to physiological processes even when the specific roles of the deformations are unknown. Development of additional tools to track organelle motions with comparable resolution as is possible with cytoskeletal filaments may be necessary for this analysis.

## ACKNOWLEDGEMENTS

We thank members of David Kovar’s laboratory, particularly J. Christensen, J. Winkelman, C. Suarez, and A. Harker for the purified protein cross-linker fimbrin, as well as the construct for fascin and advice on purifying proteins. This research was partially supported by the University of Chicago MRSEC, funded through the National Science Foundation award DMR-2011854. This research was also supported in part by the National Science Foundation EPSCoR Program under NSF Award # OIA-1655740 and the National Institutes of Health National Institute of Biomedical Imaging and Bioengineering Training grant number T32EB009412.

