## Supplemental Figures for "Direct detection of deformation modes on varying length scales in active biopolymer networks"

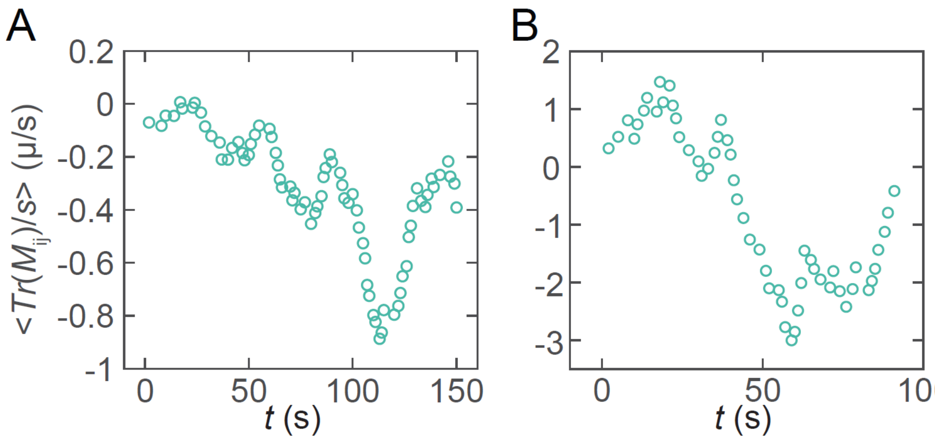


Figure S1: The contractile moment at large *s* decreases after addition of myosin. Contractile moments are plotted as a function of time for sparsely cross-linked (A) and unipolar bundle (B) networks with *s* = 50 μm.

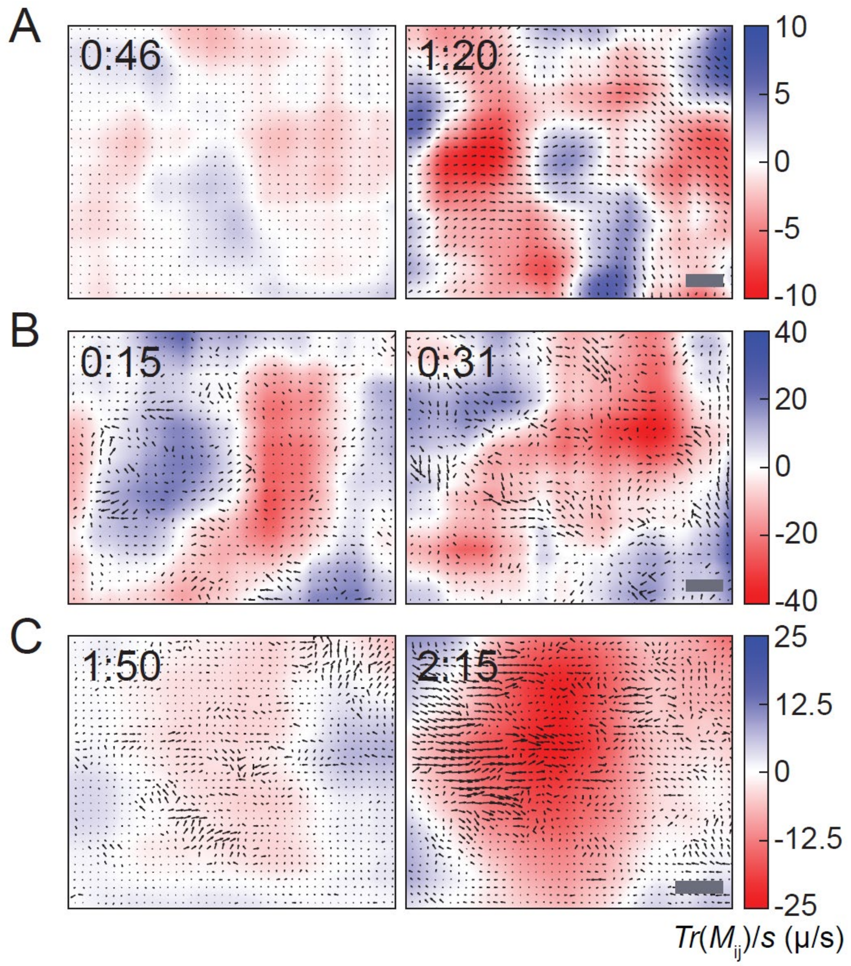


Figure S2: Large contractile regions develop early and intensify during network contraction. Colors indicate the contractile moment value for a box of *s* = 50 μm centered at the given pixel in sparsely cross-linked (A), unipolar bundle (B), and random polarity bundle (C) networks.

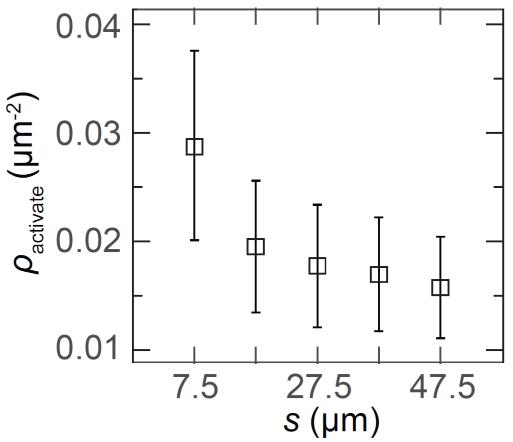


Figure S3: The myosin density required to increase $P_{biaxial}$ by 20 percent of its baseline at low $\rho_{myo}$, $\rho_{activate}$, decreases with increasing $s$ in sparsely cross-linked networks. Data points shown are the average of three to four samples.

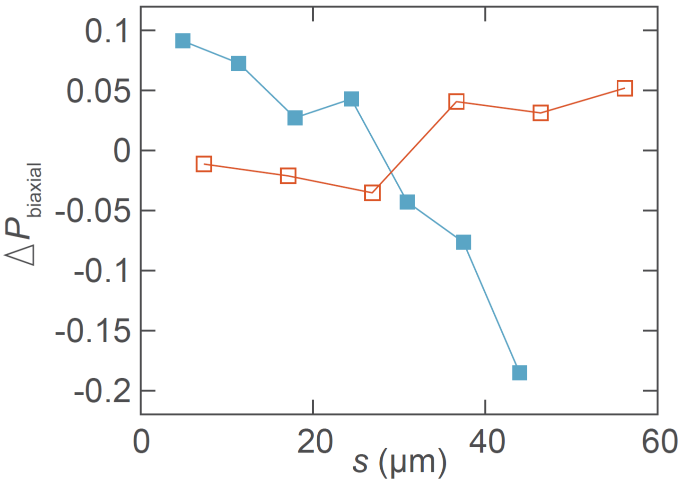


Figure S4: The change in $P_{biaxial}$ from its initial value to its extremum before the peak in $\rho_{myo}$ differs between unipolar bundle network and random polarity bundle network. Unipolar bundled data is shown with open squares and random polarity bundle data is shown with closed squares.

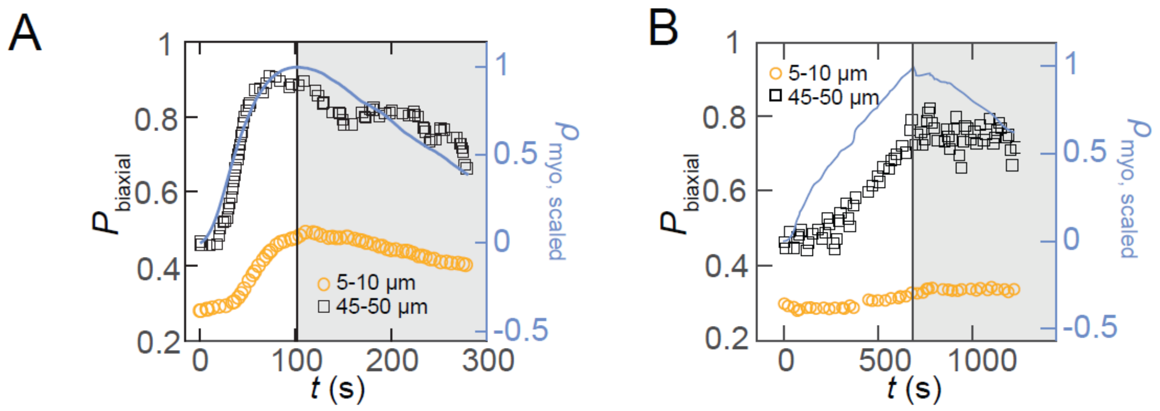


Figure S5: $P_{\mathrm{biaxial}}$ reaches its peak or plateau value at approximately the peak in $\rho_{\mathrm{myo}}$ in sparsely cross-linked networks. $P_{\mathrm{biaxial}}$ at two length scales and $\rho_{\mathrm{myo}}$ scaled between 0 and 1 are shown for (A) high $\rho_{\mathrm{myo}}$ network and (B) low $\rho_{\mathrm{myo}}$ network.
